# SHP2-triggered endothelium activation fuels estradiol-independent endometrial sterile inflammation

**DOI:** 10.1101/2024.01.16.575960

**Authors:** Jie Pan, Lixin Zhao, Wen Fang, Jiao Qu, Linhui Zhai, Minjia Tan, Qiang Xu, Qianming Du, Wen Lv, Yang Sun

## Abstract

Endometrial hyperplasia (EH) is a common gynecological disease primarily driven by excess estrogen. How endometrial sterile inflammation regulates EH remains unclear. First, we found the upregulation of SHP2 in endometrial endothelial cells from patients with EH by scRNA-Seq. SHP2 promoted inflammatory activation of endothelial cells, which promoted macrophage transendothelial migration. Subsequently, IL1β and TNFα from macrophages gave a feedforward loop to enhance endothelial cell activation and result in more IGF1 secretion, thereby sustaining sterile endometrial inflammation and facilitating endometrial epithelial cell proliferation even after estradiol withdrawal. Mechanistically, results of bulk RNA-Seq and phosphoproteomic analyses showed that endothelial SHP2 dephosphorylated RIPK1^Y380^ after estradiol stimulation. This event promoted activator protein 1 (AP-1) activation, instigating inflammation and increased CXCL10, CXCL13, COX2 and IGF1 secretion. Furthermore, targeting SHP2 by SHP099 or endothelial-specific SHP2 deletion alleviated EH progression in mice. Collectively, our findings demonstrate that SHP2 mediates the transition of endothelial activation, from estradiol-driven short inflammation to macrophage-amplified continuous sterile inflammation. Targeting chronic sterile inflammation mediated by endothelial cell activation is a promising strategy for non-hormonal intervention in EH.

## Introduction

Endometrium hyperplasia (EH) is a common uterine pathology, primarily driven by excess estrogen exposure without the counterbalancing progestin in women [1]. Estrogen stimulates cell proliferation and differentiation, whereas progesterone inhibits the effects of estrogen[2]. The predominant morphological change of EH is the thickened endometrium, while the primary histological change is the elevated endometrial gland-to-stroma ratio and cytological atypia because of irregular proliferation of the endometrial glandular epithelium[3]. It has been widely recognized that EH involves in reproductive failure and if left untreated, EH can progress to endometrial cancer[4]. About 40% of EH patients with atypia progress to develop endometrial carcinoma[5].

Progestin supplementation is the only nonsurgical management to treat EH. However, some patients exhibit no response or resistance to hormone therapy[6]. Moreover, some studies reported a significant increase in the risk of progression to carcinoma after medroxyprogesterone acetate therapy[7]. Other compounds such as genistein[8], danazol[9], metformin[10] and gonadotropin-releasing hormone therapy[11] were reported to possess therapeutic effects on EH. In addition to targeting estrogen receptor[12], aromatase inhibitors[13] were also proposed to be promising targeted strategies for EH. However, due to our limited understanding of the biological mechanisms underlying EH, there is no single compound or target for clinical treatment except progesterone. Therefore, it is essential to study the pathological mechanisms in EH. Currently, advanced high-throughput sequencing, multiomics technologies and information technologies have expanded our view of the molecular basis to a single cell level. Therefore, molecularly targeted therapies may be a promising and precise direction of EH therapy.

Abnormal epithelial cell proliferation in EH is not only regulated by hormone, but also by other factors, including cell-cell interactions, cytokines and growth factors[1, 14]. It has been reported that the infiltration of immune cells increases as the disease progresses through different stages[2]. Besides, the close association between EH and natural killer cells, T cells, macrophages and neutrophils has been confirmed[1, 15]. In particular, macrophages are considered to play key roles in many gynecological diseases[16, 17]. Therefore, tissue inflammation may be the primary regulators of EH. However, there is no clinical applications of anti-inflammation drugs or strategies that targeting molecules in inflammation in EH treatment. Insufficient understanding underlying the molecular mechanism of tissue inflammation function in EH is the most likely reason.

Endothelial cells are the most important regulators of organ-specific immune response and tissue inflammation[18]. Almost all pathological events of inflammation are associated with the dysfunction of endothelial cells. The activated endothelial cells lead to blood flow, vascular permeability and leukocytes transendothelial migration to tissue[19, 20]. Besides, activated endothelial cells function as inflammatory cells that secreting many extracellular proteins, including chemokines, cytokines and growth factors[19]. However, most studies on inflammation focused mainly on immune cells and parenchymal cells, rather than endothelial cells. The mechanisms behind endothelial cell activation in inflammation and targeted therapy for these processes remain limited. Therefore, investigating new mechanisms of endothelial cell activation and identifying therapeutic targets offer promising approaches to control tissue inflammation and thus mitigating inflammation in EH development.

The effects of protein kinases have been well studied in EH[21–25]. However, the function of protein tyrosine phosphatase in EH progression is previous obscure. SHP2, a famous protein tyrosine phosphatase encoded by *PTPN11*, has been considered as a target in many inflammatory diseases. SHP2 regulates inflammation through regulating the phosphorylation of tyrosine on its substrate protein[26–28]. SHP2 is a positive regulator for receptor protein tyrosine kinases in mediating cellular response to hormones[29, 30], growth factors[31], and cytokines[32]. As a hot therapeutic target, clinical trials have been conducted on many SHP2 inhibitors[33]. The function of SHP2 in human reproduction is now beginning to be uncovered. SHP2 is not only involved in the pathologenesis of female infertility[34] and endometriosis[35], but also plays essential roles in the physiological function of uterus such as embryo implantation and stromal decidualization[36]. The importance of SHP2 in endometrium indicates that SHP2 possesses potential roles in EH. However, whether and how SHP2 participates in the pathogenesis of EH remains unknown.

In this study, we elucidate that SHP2 in endothelial cells responses to excess estradiol stimulation and thus drives inflammatory activation of endothelial cells. This response is achieved through the dephosphorylation of RIPK1 at Y380 site by activating the transcript factor complex AP-1, which in turn promotes inflammatory gene transcription. Irrespective of initial estrogen-induced inflammation, the activated endothelial cells secret IGF1 and inflammatory mediators to engender and maintain a long-term endometrium inflammation through recruiting macrophages. Subsequently, the cytokines from macrophages reshape the inflamed endometrium and further enhance the inflammatory activation of endothelial cells by a positive feedback loop. Increased IGF1 released by inflammatory cells leads to epithelial cell proliferation. Collectively, the results indicate the importance of activated endothelial cells and continuous estradiol-independent sterile inflammation as regulators of EH. Targeting tissue inflammation through controlling endothelial activation by SHP2 inhibition or other anti-inflammation targets such as RIPK1 may be potential interventions for EH.

## Results

### 1. Endothelial SHP2 is abnormally increased and associated with EH pathogenesis

We have been interested in the function of protein tyrosine phosphatases (PTPs) in diverse chronic inflammatory diseases for a long time[26, 37]. To investigate the role of PTPs in EH, we first collected 11 normal endometrium samples and 12 EH endometrium samples for single cell RNA sequencing (scRNA-Seq) to profile all the PTPs (**Fig.1A**). We identified 15 major clusters of different cell types (**Supplementary Fig. S1A**). To study different PTPs expression between normal and EH samples, we analyzed all detected genes encoding PTPs in 15 cell types (**Supplementary Fig. S1B**). Unexpectedly, there were few different expressed genes in epithelial cells but in endothelial cells between normal and EH group. The results showed that there were 4 genes (*DUSP23*, *DUSP11*, *DUSP6* and *PTPN11*) with high expression and significantly different expression between normal and EH in endothelial cells (**Fig.1B**). Next, we found the mRNA expression of *PTPN11* which encodes the protein tyrosine phosphatase SHP2, was the highest among the 4 different expressed genes in endothelial cells (**Fig.1C**). Besides, *PTPN11* was the specifically changed gene in endothelial cells but not in other cell types between normal and EH samples (**Fig.1D**). Thus, we hypothesized that the increased *PTPN11* expression in endothelial cells of EH tissues may play a part in EH pathogenesis.

**Figure 1.**
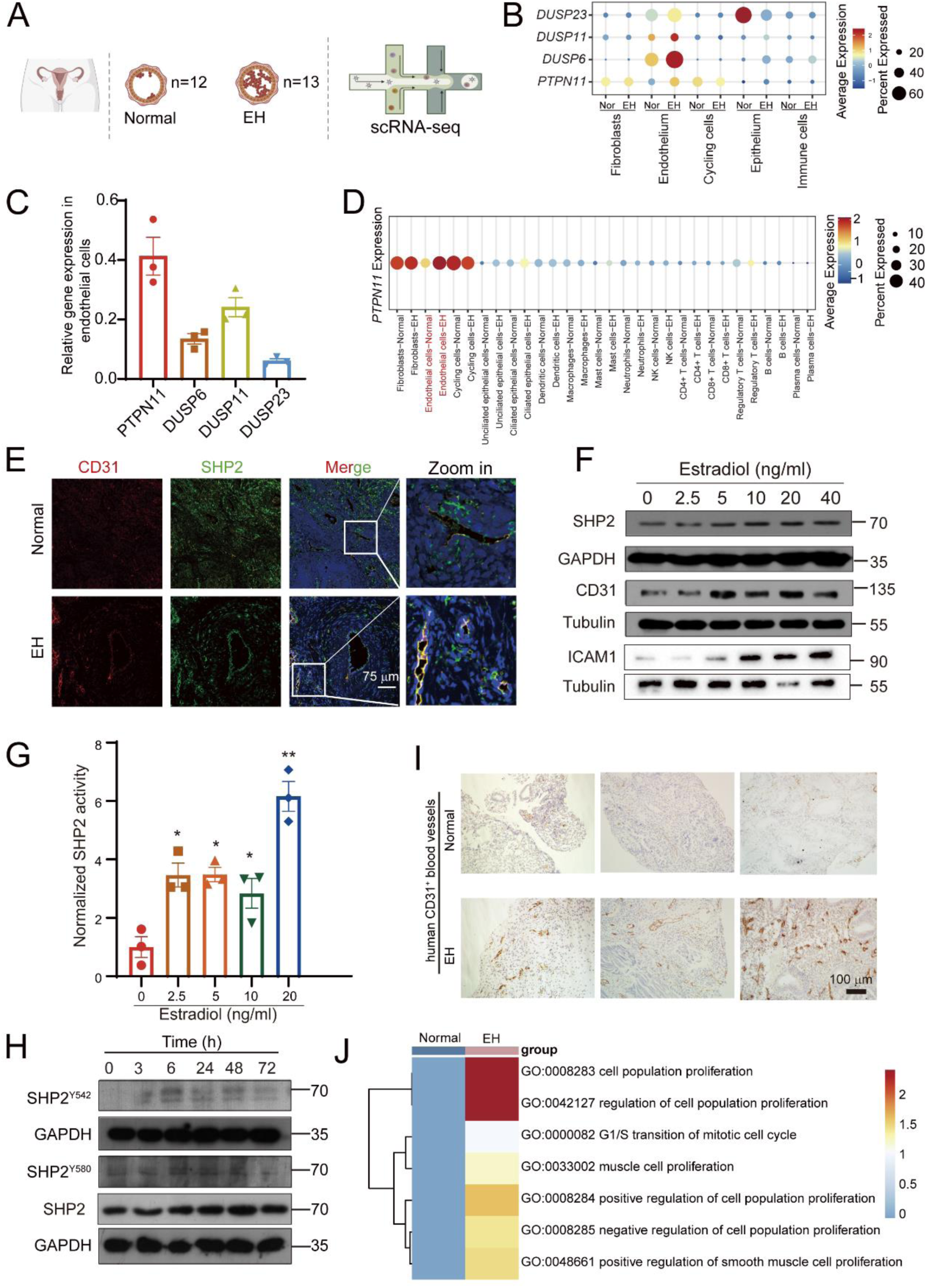
SHP2 in endothelial cells was upregulated and activated by estradiol in endometrial hyperplasia tissue. (**A**) Schematic representation of clinical human endometrium samples collection for single-cell RNA sequence (scRNA-Seq). Normal represents healthy normal endometrium, n=12; EH represents endometrium samples from endometrial hyperplasia patients, n=13 (**B**) Dot plot showing the expression of 4 kinds of protein tyrosine phosphatases in different kinds of cell types in human endometrium tissue from the scRNA-Seq data of clinical samples. Nor represents normal endometrium samples. EH represents EH endometrium samples. (**C**) Relative mRNA expression of the 4 indicated genes in HUVECs, determined by real-time quantitative polymerase chain reaction (qPCR). (**D**) Dot plot showing the different expression of *PTPN11* in different cell types from samples of normal endometrium (Normal) and EH endometrium (EH). (**E**) Expression of SHP2 (green) in CD31^+^(red) vessels from samples of normal endometrium and EH endometrium (n=5, respectively). Scale bar = 100 μm (**F**) The protein expression of SHP2, CD31 and ICAM1 was examined in HUVECs following treatment with different doses of estradiol for 24 h. (**G**) SHP2 enzyme activity was assessed in HUVECs after treated with different dose of estradiol for 24 h. *p value <0.05; **p value <0.01 compared with the control group (0). (**H**) IHC staining of CD31^+^ vessels in samples of normal endometrium and EH endometrium (n=5, respectively). (**I**) The protein of phosphorylated SHP2 in HUVECs treated with 40 ng/mL estradiol for indicated time. (**J**) GSEA analysis in endothelial cells from the scRNA-Seq data of clinical endometrium samples. All results are representative of two or more replicates. Data were analyzed by Dunnett *t* test in Figure 1G.

To confirm the protein expression of SHP2 in clinical samples, we conducted immunofluorescence staining. The results showed that the expression of SHP2 was increased in EH endometrial vessels (labeled by CD31, red) (**Fig. 1E**). Based on these results, we wondered whether the increased SHP2 expression was directly regulated by estradiol stimulation. Thus, we cultured human umbilical vein endothelial cells (HUVECs) with different doses of estradiol for 24 h. As we expected, the expression of SHP2 in HUVECs was upregulated by estradiol **(Fig.1F & Supplementary Fig. S1C)**. The enzyme activity of SHP2 increased after estradiol stimulation (**Fig.1G**). The phosphorylation of Y542 and Y580 are indicators of SHP2 activation. Our data also demonstrated that the increased activity of SHP2 may be attributed to its increased phosphorylation at Y542 and Y580 (**Fig. 1H**). However, consistent with the results of scRNA-Seq, SHP2 expression was not influenced by estradiol in human endometrium epithelial cells (hEECs). It was possible because of lower basal SHP2 expression in hEECs compared with HUVECs (**Supplementary Fig. S1D, E**). Besides, CD31 and ICAM1, markers of endothelial cell activation, were increased after estradiol treatment (**Fig.1F, G**). EH often accompanies angiogenesis and abnormal bleeding, which is a result of endothelial cell dysfunction[38, 39]. Thus, we next detected the state of the endometrial tissue vasculature. We found that the endometrial vessel number increased in EH samples. Alongside, scRNA-Seq analysis revealed the upregulation of the proliferation signaling pathway in endothelial cells (**Fig.1 I, J**). Hence, our results identified that the expression and enzyme activity of SHP2 were significantly increased in endothelial cells by excessive estradiol stimulation, leading to vascular dysfunction in EH.

### 2. Endothelial specific deletion of SHP2 alleviates estradiol-induced EH in mice

To investigate the direct role of endothelial SHP2 in EH pathogenesis, we obtained transgenic mice of inducible endothelial-specific SHP2 knockout mice (SHP2^iECKO^, **Supplementary Fig. S2A, B**). Seven days after the final tamoxifen injection, we constructed EH model by exposure to estradiol for 21 consecutive days. The treatment did not influence the body weight of mice (**Supplementary Fig. S2C**). The results of immunofluorescence staining and WB of lung endothelial cells showed that SHP2 expression decreased in endothelial cells following 5 days of tamoxifen injection (**Supplementary Fig. S2D-F**). Compared to the WT vehicle control mice, the uterus of mice from other 3 groups were markedly larger. While the uterus from SHP2^iECKO^ mice were smaller than that from SHP2^f/f^ mice (**Fig.2A**). The uterine weight of SHP2^iECKO^ mice was significantly decreased compared to that of SHP2^f/f^ mice (**Fig.2B**). Histological evaluation of the mouse uterus after estradiol exposure revealed a proliferative endometrium with architectural abnormalities and increased gland-to-stroma ratio in both WT and SHP2^iECKO^ group (**Fig.2C, D**). However, the histological changes of EH were slight and relieved in SHP2^iECKO^ mice (**Fig.2C**). We also observed that the number of endometrial vessels was increased in the WT and SHP2^f/f^ groups after estradiol exposure, while it was less in the SHP2^iECKO^ group (**Fig.2E**). All our data demonstrated that endothelial SHP2 was positively involved in EH progression.

**Figure 2.**
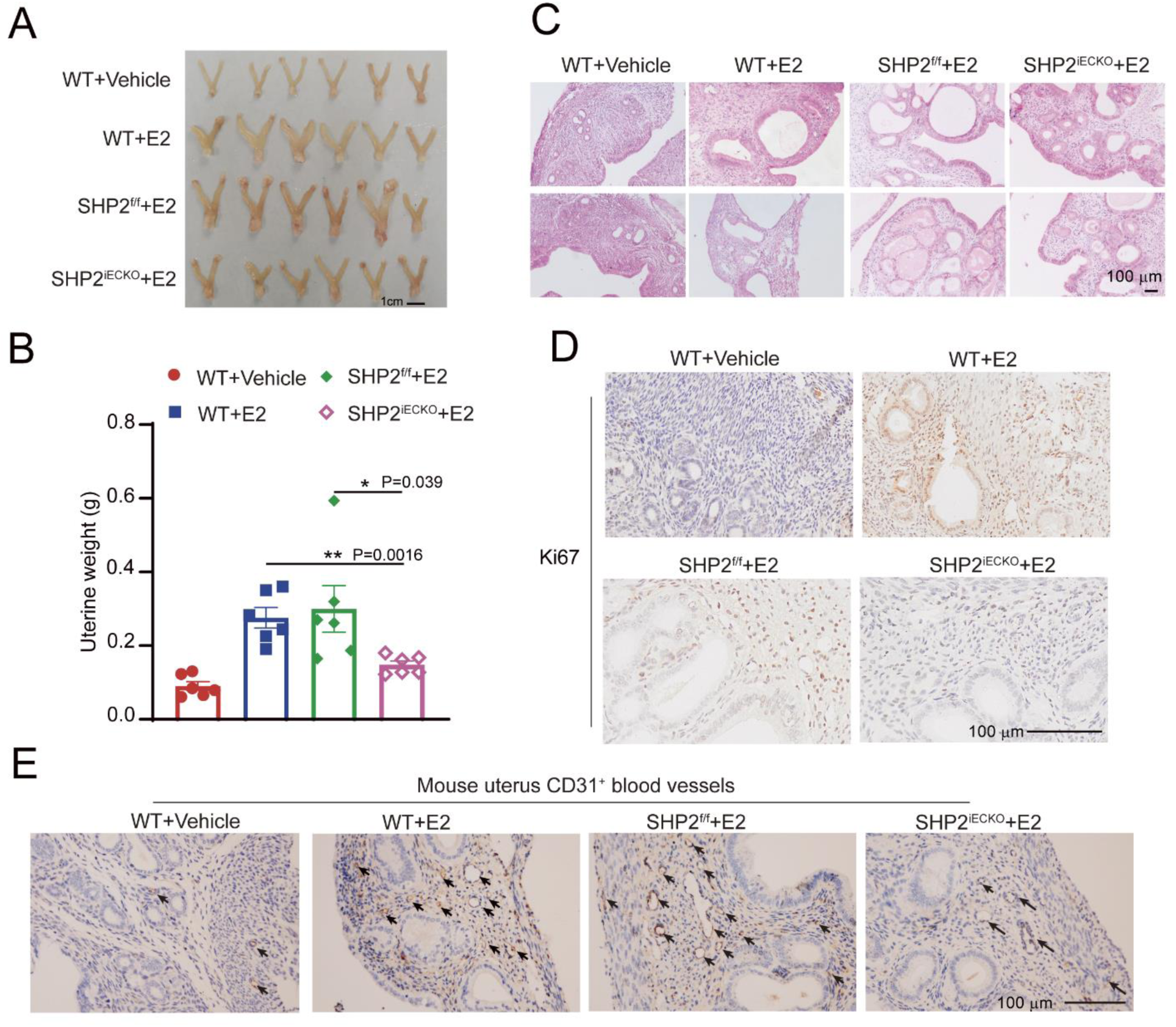
SHP2 specific deletion in endothelial cells reversed estradiol induced EH in mice. (**A**)The morphology of mice uterus from 4 different groups of mice with indicated treatment (n=6/group). (**B**) H&E staining of mice uterus from the 4 groups (n=6/ group). (**C**) The uterine weight of mice from the indicated group (n=6/group). (**D**) IHC staining of Ki67^+^ proliferative cells in the mice uterus from the 4 groups. Body weight curve of mice from indicated group (**E**) IHC staining of CD31^+^ vessels in the mice uterus from the 4 groups. WT+ Vehicle group represents the WT C57BL/6 mice treated with olive oil for 21 days. WT+E2 group represents the WT C57BL/6 mice treated with estradiol (100 μg/kg) for 21 days. SHP2^f/f^ +E2 group represents the *Shp2*^flox/flox^ transgenic mice with C57BL/6 background treated with estradiol (100 μg/kg) for 21 days. SHP2^iECKO^ +E2 group represents the specific endothelial cells SHP2 deletion transgenic mice with C57BL/6 background treated with estradiol (100 μg/kg) for 21 days. Data were analyzed by Tukey-Kramer in Figure 2B.*p value <0.05; **p value <0.05.

### 3. SHP2-activated inflammatory endothelial cells significantly secret more extracellular factors

Endothelial cells can respond to locally derived signals, including estrogen[40] and transform from the resting state to the activation state[19]. Estrogen can activate GPCR to induce type I activation of endothelial cells. Type Ⅱ activation usually goes after type I activation [19]. We next characterized the molecular and functional heterogeneity of endometrium endothelial cells in healthy individuals and EH patients at the single-cell level. We subtyped all of the endothelial cells into 8 clusters (**Supplementary Fig. S3A**), and we found the endo1 subtype expressed high levels of inflammatory cytokines and leukocyte adhesion molecules that represented endothelial activation and mediated leukocyte entry, such as ICAM1, IL33, IL6, SELE and SNCG (**Supplementary Fig. S3A**). Interestingly, the inflammatory endothelial cell clusters 1 also expressed more *PTPN11* in EH samples (**Supplementary Fig. S3B**). Intracellular Ca^2+^ level is an early step of endothelial activation. Thus, we measured the Ca^2+^ level after estradiol stimulation in the presence and absence of SHP099. The results showed that estradiol treatment increased intracellular Ca^2+^ in a short time but the SHP099 pretreatment did not influence the Ca^2+^ influx in HUVECs, which indicated that the early event of type I endothelial cell activation is not dependent on SHP2 (**Supplementary Fig. S3D-G**). We observed that SHP2 overexpression increased the expression of ICAM1 and COX2, which were markers of endothelial cell activation (**Supplementary Fig. S3H-I**). Therefore, these results indicated SHP2 was a new molecule in regulating inflammatory activation of endothelial cells. Activated endothelial cells are kinds of inflammatory secretory cells. They can release extracellular proteins to remodel tissue microenvironment. To this end, we analyzed the extracellular proteins and found that the total secreted proteins were significantly increased in endothelial cells from EH patients detected by scRNA-Seq (**Fig. 3A-B**). To test whether the extracellular proteins found in scRNA-Seq data also increased in activated endothelial cells by estradiol stimulation, we further detected the extracellular proteins closely associated with inflammation and proliferation, including CXCL10, CXCL13, PTX3, WNT5A, FRZB, OSM and IGF1 in HUVECs after estradiol stimulation (**Supplementary Fig. S3J**). The results showed that SHP2 induced CXCL10, CXCL13 and IGF1, expressed in the activated endothelial cells (**Fig. 3C-D**). IGF1 is an essential growth factor which has been reported to be involved in the proliferation of endometrial glandular epithelium[41]. Endothelial expression of CXCL10, CXCL13 and IGF1 were reversed by inhibition of SHP2 with SHP099 (**Fig. 3F-G**). To confirm the endothelial activation in tissue, we performed immunofluorescence staining of mice uterus and identified the increase of activated endothelial cells (ICAM1^+^CD31^+^) in EH mice compared with vehicle control and SHP2^iECKO^ mice (**Fig. 3E**). We also observed increased IGF1 expression in EH mice uterus and especially increased IGF1 staining around blood vessel structure (**Fig. 3F**). However, IGF1 expression decreased in the uterus of SHP2^iECKO^ mice, which suggested IGF1 was positively regulated by endothelial SHP2 (**Fig. 3F**). Next, we also observed SHP2 mediated endothelial inflammatory activation in HUVECs indicated by high levels of ICAM1 and VCAM1 after estradiol stimulation (**Fig. 3G**). Together, our data suggest that SHP2 triggered the endothelial activation and promoted CXCL10, CXCL13 and IGF1 secretion after estradiol stimulation.

**Figure 3.**
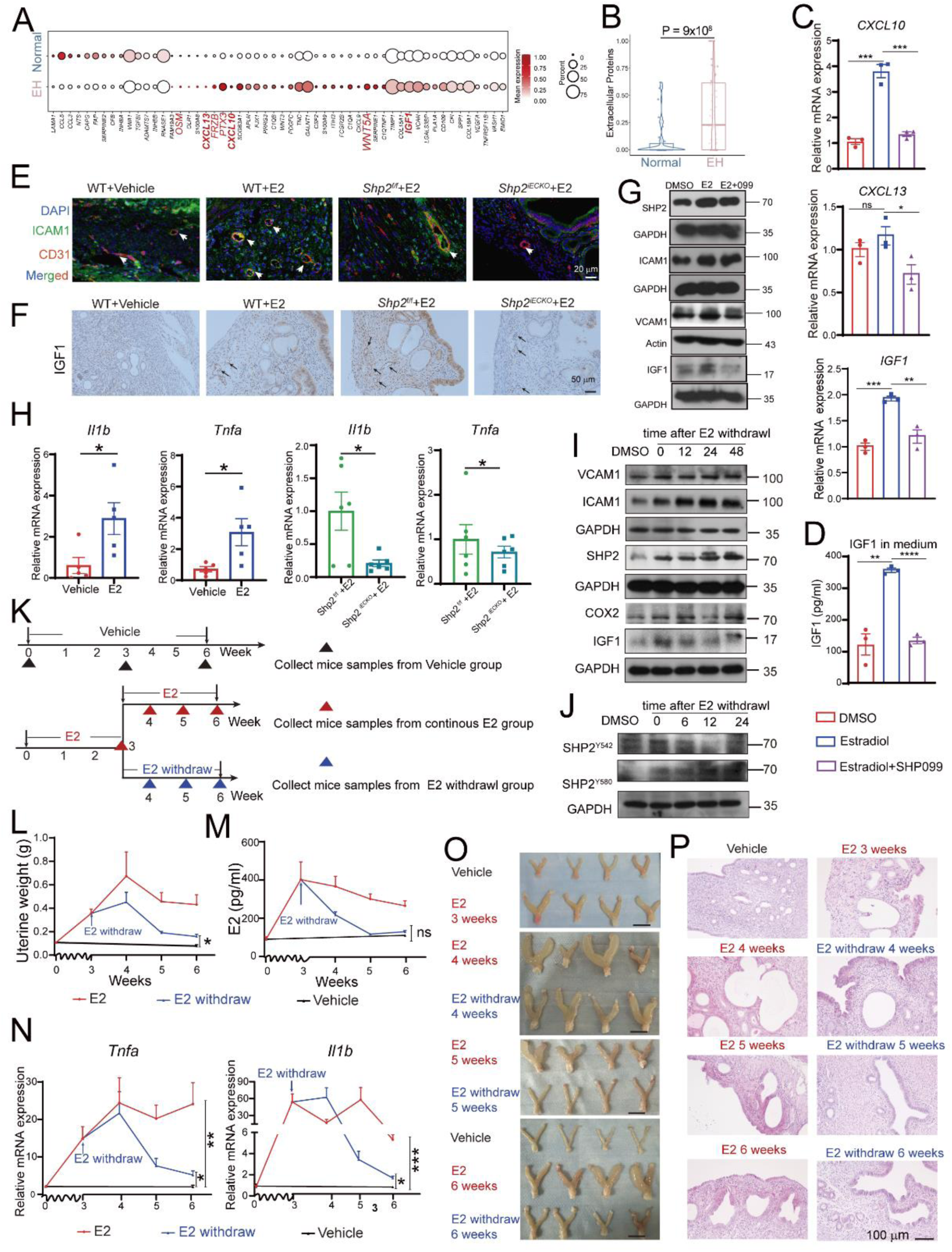
Endothelial SHP2 promoted endothelial inflammatory activation and sustained a long-term sterile tissue inflammation. **A**, Dot plot showing the expression of differently expressed genes of secreted extracellular proteins in inflammatory activated endothelial cells from scRNA-Seq data. **B**, Statistic analysis of all the extracellular protein level in endothelial cells from scRNA-Seq data of healthy normal endometrium and EH endometrium. **C**, Validation of the indicated extracellular proteins in HUVECs by qPCR after stimulated with 40 ng/mL estradiol (E2) in the presence and absence of SHP099 (5 μM) for 24 h. **D**, IGF1 level in HUVECs medium detected by Elisa after stimulated with 40 ng/mL estradiol (E2) in the presence and absence of SHP099 (5 μM) for 48 h. **E**, Detection of inflammatory activation of endothelial cells (CD31^+^ICAM1^+^) in mice uterus with indicated treatment (n=6/group). **F**, Tissue IGF1 expression by endothelial cells by IHC staining in mice uterus tissue from 4 indicated groups (n=6/group). **G**, WB analysis of the proteins that indicated endothelial cell inflammatory activation in HUVECs treated with estradiol (E2) for 24 h and then withdrew estradiol (E2) to left the HUVECs untreated for indicated time. **H**, The mRNA expression of *Il1b* and *Tnfa* in mice uterus tissues treated with estradiol (E2) or vehicle in WT or SHP2^f/f^ or SHP2^iECKO^ mice. n=5 in WT group, n=6 in SHP2^f/f^ group, n=6 in SHP2^iECKO^. **I**, WB analysis of the proteins that indicated endothelial cell inflammatory activation in HUVECs treated with estradiol (E2) for 24 h and withdrew estradiol (E2) to left the HUVECs for indicated time. **J**, WB analysis of phosphorylation of SHP2 after estradiol treatment for 3 h and then withdrew estradiol (E2) to left the HUVECs untreated for indicated time. **K**, Design of the animal experiment to determine the role of endothelial activation-mediated tissue inflammation in EH after estradiol (E2) withdrawal. Mice uterus from vehicle and E2 group were first collected at 3 weeks to confirm the success of EH mouse model. Mice from E2 group were then divided into E2 and E2 withdrawal group, respectively. E2 withdrawal group was first treated with estradiol for 3 weeks to generate EH model and then left untreated for another 3 weeks. Mice from E2 group were treated with continuous estradiol for 6 weeks. Thereafter, the mice uterus was collected at 4, 5 weeks from E2 and E2 withdrawal group. Finally, at 6 weeks, mice uterus from vehicle, E2 and E2 withdrawal groups were collected. **L**, Uterine weight from indicated groups. **M**, E2 concentration in serum from mice of indicated group detected by Elisa. **N**, The mRNA expression of *Tnfa* and *Il1b* in mice uterus at different time point. Group vehicle, n=8; group E2 at 3 weeks, n=4; group E2 at 4 weeks, n=4; group E2 at 5 weeks, n=5; group E2 at 6 weeks, n=5; group E2 withdraw at 4 weeks, n=6; group E2 withdraw at 5 weeks, n=7; group E2 withdraw at 5 weeks, n=6. **O**, The morphology of mice uterus from indicated group at indicated time point. Scale bar: 1cm. **P**, H&E staining of the histological change of mice uterus from indicated group at indicated time point. Data are represented as mean ± SEM. Data were analyzed by Student’s *t* test in Figure 3B and H, and Tukey-Kramer in Figure 3C and D, and Dunnett test in Figure 3L, M and N. *p value <0.05; **p value < 0.01, ***p value <0.001; ns represents no significant difference. See also figure S3 and figure S4. **p value < 0.01; ***p value <0.001; ****p value < 0.0001; ns represents no significant difference.

### 4. SHP2-triggered endothelial cell activation generates and sustains estradiol independent sterile inflammation in EH

Activation of vascular endothelial cells is involved in tissue sterile inflammation[42]. Previous studies have reported that neutrophils and T cells are significantly increased in EH patients[17]. In addition to being driven by estrogen, endometrial proliferative activity cane promoted by cytokines and growth factors secreted by cells in the stroma[2]. Therefore, we proposed that endothelial cells activation generated endometrium inflammatory environment that involved in the progression of EH. Based on the hypothesis, we first analyzed the expressions of *Il1β* and *Tnfα* were significantly increased in the uterus tissues of EH mice (**Fig. 3H**). While the level of *Il6* remained unchanged in EH mice group compared to the vehicle control (**Supplementary Fig. S3K**). Moreover, these inflammatory cytokines decreased in the uterus of SHP2^iECKO^ mice compared to the SHP2^f/f^ group **(Fig. 3H)**. The results indicated that estradiol induced uterine inflammation was mediated by endothelial SHP2. Consistently, COX2 expression was also upregulated in the endometrium samples of EH patients and EH mice uterus (**Supplementary Fig. S3L and Fig. S4A).** Hence, all these results showed that tissue sterile inflammation existed in the uterus of EH patients or in the uterus of EH mice. The uterine inflammation was dependent on the expression of SHP2 in endothelial cells.

Because estrogen level is not that high all the time in EH diseases, we therefore asked whether and how the existed tissue inflammation in the endometrium could sustain and promote the progression of EH. We first treated HUVECs with estradiol for 48 h and then withdrew estradiol at various intervals, the results showed a sustained expression and activation of SHP2, as well as high levels of ICAM1, VCAM1, COX2 and IGF1 (**Fig. 3I-J**). To validate sterile inflammation independent of continuous estradiol exposure *in vivo*, we constructed EH mouse model by 3 weeks of estradiol injection, and then 20 mice from estradiol treated group were then withdrawn from estradiol for one to three weeks, and the remaining mice (EH group) continued to be exposed to estradiol (**Fig. 3K**). We collected the uterus and serum of mice from the 3 groups at different time points to observe the progression of EH and tissue inflammation (**Fig. 3L-N**). The results showed that mice uterine weight and size in sustained 6 weeks-estradiol exposure group increased with time (**Fig. 3L and 3O**). In the estradiol withdrawal group, serum estradiol levels quickly decreased and recovered to baseline after 2 weeks estradiol withdrawal (at the time point of 5 weeks) (**Fig. 3M**). However, the mice uterine weight and size in estradiol withdrawal group also continued to increase even after being left untreated for a week (at the time point of 4 weeks), and then fall back after 2 or 3 weeks after estradiol withdrawal (at the time point of 5 weeks and 6 weeks). Interestingly, the absolute weight of mice uterus in estradiol withdrawal group was still significantly higher than that of vehicle control (**Fig. 3L and 3O**). The EH phenotype of endometrium still existed and did not recover to the normal state. In line with this, the tissue showed persistent histological changes and sustained inflammation (**Fig. 3P and Supplementary Fig. S4B, C**). Moreover, the expression of tissue IGF1 and endothelial SHP2 (CD31^+^ SHP2^+^ endothelial cells) was keeping at high level even after estradiol withdrawal (**Supplementary Fig. S4D, E**). All these results indicated that after the construction of EH model at 3 weeks, the tissue inflammation environment and endothelial activation in mice uterus sustained at high level. This persisted even after the estradiol fell down to normal levels, suggesting that tissue could not recover to a normal state. In summary, the high expression of SHP2-mediated endothelial cells activation generated and sustained chronic tissue inflammation in uterus of EH mice. The long-term endometrium inflammation was also involved in EH pathology. Exposure to estradiol is not necessary for the long-term sterile inflammation, but it is the driver of endometrium inflammation and EH. There may be some other factors responsible for continuous amplification of endometrial inflammation.

### 5. Inflammatory endothelial cells sequentially traffic and activate macrophages to amplify inflammation

Almost all the pathological events of inflammation are associated with the dysfunction of endothelial cells through the increased blood flow, vascular permeability and transendothelial migration of leukocytes [20]. Therefore, we next examined what kind of immune cells endothelial cells traffic into endometrium of EH. We characterized the network of interaction between endothelial cell-immune cell by scRNA-seq data analysis (**Supplementary Fig.S5A, B**). The interactions between endothelial cell and macrophages were significantly enhanced in EH samples **(Fig. 4A)**. Besides, the cell number and proportion of endometrial macrophages in EH samples were higher than other immune cells, accounting for 62% **(Fig. 4B)**. We observed substantial numbers of infiltrating macrophages (F4/80^+^) frequently localized closely to CD31^+^ blood vessels (**(Fig. 4C).** In line with the findings, the IHC staining of F4/80^+^ macrophages were increased in EH mice uterus and decreased in SHP2^iECKO^ mice, which suggested the infiltration of macrophages was dependent on the endothelial SHP2 expression (**Fig. 4D**). To address the interaction between endothelial cells and macrophages, we first treated HUVECs with estradiol for 48 h and replaced the medium to coculture HUVECs and THP1 cells for another 1 h. The estradiol-activated endothelial cells effectively increased the adhesion of THP1 cells to HUVECs. (**Fig. 4E, F**). Contrary to our hypothesis, estradiol could not directly induce THP1 activation (**Supplementary Fig. S5C**) but the conditional medium (CM) from estradiol treated endothelial cell promoted macrophage activation and released IL1β, TNF and IL6 (**Fig. 4G, H**). While SHP099 reversed the effect of estradiol cultured HUVECs CM on the activation of macrophage. We also used the CM from SHP2 overexpressed HUVECs to treat THP1 cells, it had stronger effect on macrophages activation compared with that of estradiol treated HUVECs CM (**Supplementary Fig. S5D**). All of these results demonstrated that high expression of SHP2 induced by estradiol promoted endothelial activation, which then attracted, activated macrophages, and ultimately generated an inflammatory endometrium environment. We cultured HUEVCs with CM from THP1 cells pretreated with or without PMA for 24 h (**Fig. 4I**). The results showed that CM from activated macrophages upregulated the protein expression of SHP2, ICAM1, VCAM1, COX2 and IGF1 (**Fig. 4J**). Consistent with the increased protein level, the mRNA level of *PTPN11, PTGS2* and the adhenrens and junction molecules, including *ICAM1* and *SELE*, increased (**Fig. 4K**). The inflammatory cytokines and chemokines, including *CXCL10*, *CXCL13*, *IL6* and *IL33*, also increased in HUVECs after treating with CM from activated THP1(**Fig. 4L**). Moreover, the mRNA expression and protein expression of IGF1 in HUVECs culture supernatant were significantly upregulated in HUVECs by treatment of CM from activated THP1 (**Fig. 4L-M**)). To mimic the function of HUVECs CM in activated macrophages, we stimulated HUVECs with estradiol or with different kinds of inflammatory cytokines. The results showed that the effects of inflammatory factors on SHP2 expression and endothelial activation were stronger than the estradiol stimulation alone (**Supplementary Fig. S5E**). Together, these data uncovered that excessive estradiol initially activated endothelial cells through upregulating SHP2, and then the activated endothelial cells facilitated tissue macrophages which reshape the inflammatory uterine environment. Through a feedforward loop, the activated macrophages further amplified endothelial inflammatory activation through cytokines such as IL1β and TNFα to sustain the chronic sterile endometrial inflammation and to secret more growth factor IGF1.

**Figure 4.**
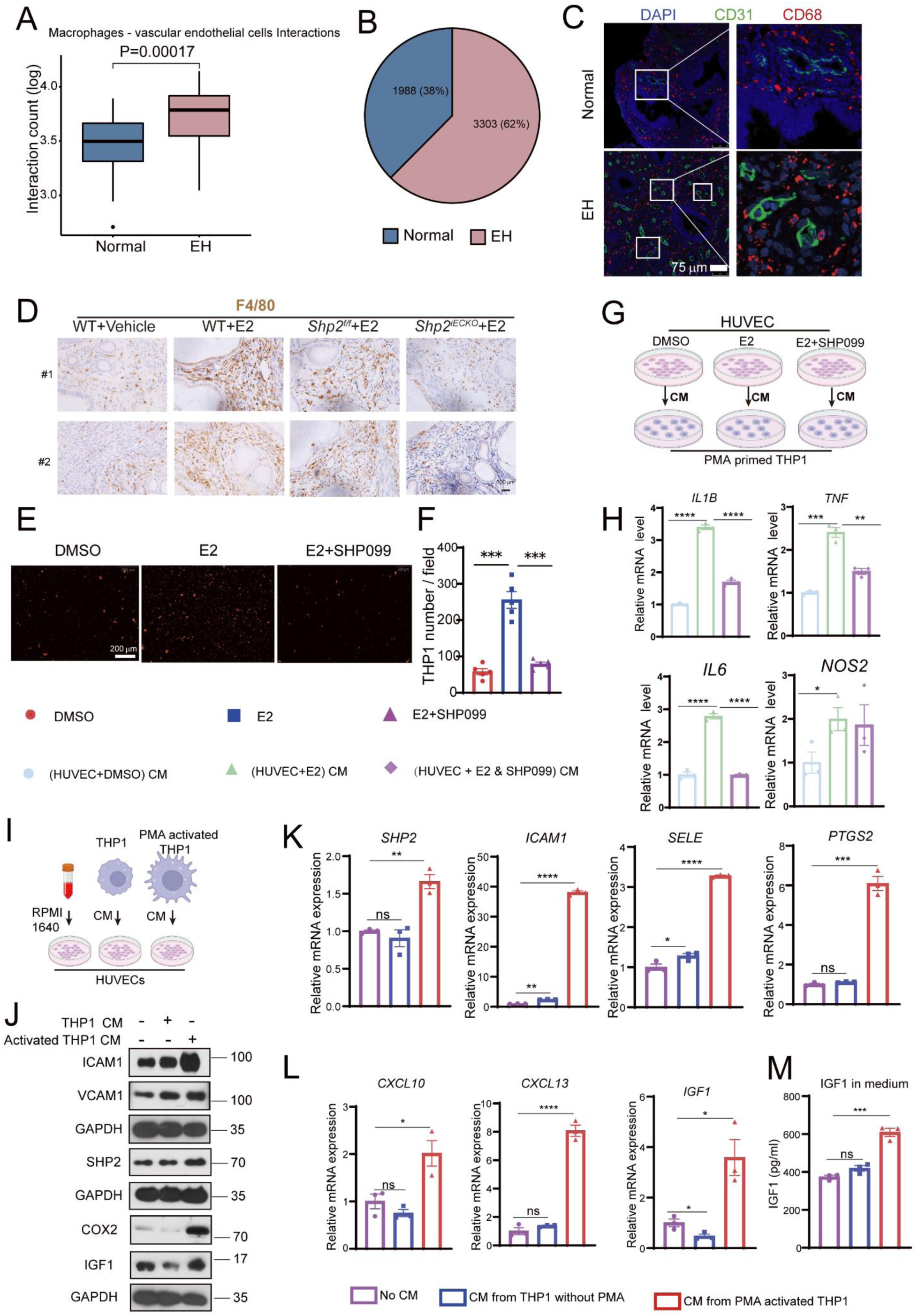
Activated endothelial cell - macrophages crosstalk got positively fed back to endometrial sterile inflammation. **A**, Interactions between macrophages and vascular endothelial cells predicted by CellPhoneDB. **B**, Pie chart showing the cell number and percentage of macrophages in Normal and EH group, respectively. **C**, Immunofluorescence analysis of endothelial cells (CD31, green), macrophages (F4/80, red) in endometrium from Normal or EH patients. **D**, IHC staining of F4/80^+^ macrophages in mice uterus tissues treated with estradiol (E2) or vehicle in WT or SHP2^f/f^ or SHP2^iECKO^ mice. **E**, The attachment of Dil labeled THP1 cells to HUVECs after estradiol (E2, 40 ng/mL) with or without SHP099 (5 μM) treatment for 24 h. **F**, Statistic analysis of the number of attached THP1 cells in 5 fields of one representative experiment. **G** and **H**, QPCR detection of mRNA level of *IL1B*, *TNF*, *IL6* and *NOS2* in THP1 cells cultured with condition medium from HUVECs pretreated with DMSO or estradiol (E2, 40 ng/mL) with or without SHP099 (5 μM) for 48 h. **I**, Experimental outline of different treatment to indicated cells. The condition medium (CM) from THP1 cells treated with or without PMA (100 ng/mL) for 24 h and then replaced with complete RPMI 1640 medium for another 24 h culture. The CM from THP1 or complete RPMI 1640 was subjected to treat HUVECs for 24 h. **J**, WB analysis of proteins that indicated endothelial inflammatory activation after treatment of CM from THP1 cells. **K**, QPCR detection of mRNA level of SHP2 and the genes indicated endothelial cells inflammatory activation after CM treatment from THP1. **L**, QPCR detection of mRNA levels of genes encoding secreted extracellular proteins in activated HUVECs after CM treatment from THP1 cells. **M**, IGF1 in HUVECs medium after CM treatment from THP1. Data are represented as mean ± SEM. Data were analyzed by Tukey-Kramer. *p value <0.05; **p value < 0.01; ***p value <0.001; ns represents no significant difference. See also figure S5.

### 6. IGF1 derived from activated endothelial cells supports endometrial epithelial cell proliferation

One of the pathologic mechanisms of EH is the abnormal proliferation of endometrial epithelial cells caused by an imbalance of estrogen and progesterone 17. Our data showed that endothelial cells secreted high levels of IGF1. In light of this, we next aimed to detect the role of IGF1 and the function of activated HUVECs on the proliferation of human endometrial epithelial cells (hEECs). We first detected the effect of estradiol alone on the proliferation of hEECs. However, the estradiol failed to promote hEECs proliferation even at high concentration for 48 h (**Fig. 5A**) and had no effect on the cell cycle progression (**Supplementary Fig. S6A, B**). The results indicated that excess estradiol alone was not sufficient to initiate the abnormal proliferation of epithelial cells. There may be some other factors from the interacted cells in tissue to interact with epithelial cells and act synergistically with estrogen to promote abnormal epithelial proliferation. We had found that activated endothelial cells secreted increased IGF1. Whether this paracrine secretion contributes to abnormal proliferation of epithelial cells has not been investigated. So, we treated the hEECs with different concentrations of IGF1, the result showed that IGF1 promoted the hEECs proliferation in a dose-dependent manner (**Fig. 5B**). In line with this, the Edu assay also confirmed the positive effect of IGF1 on the proliferation of hEECs (**Fig. 5C, D**). Therefore, epithelial cells may interact with endothelial cells through IGF1 signaling, leading to abnormal epithelial cells proliferation. We also observed a significant enhancement in the interaction of endothelial cells and epithelial cells, as shown by the bule bolder line that connected endothelial cell and epithelial cell in our scRNA-Seq data (**Fig. 5E, F**). To confirm the enhanced interaction between endothelial cells and epithelial cells, we used the CM from estradiol pretreated HUVECs or THP1 CM pretreated HUVECs to further culture hEECs for 48 h. The Edu assay demonstrated that CM from estradiol or THP1 CM activated HUVECs significantly promoted proliferation of hEECs (**Fig. 5G-J**). Consistently, the CCK8 assay also showed that the CM from HUVECs had the similar effect on the proliferation of hEECs (**Supplementary Fig. S6C, D**). Overall, activated endothelial cells promoted abnormal epithelial cells proliferation through a paracrine secretion way, and one of the effector factors was the increased secretion of IGF1 from endothelial cells.

**Figure 5.**
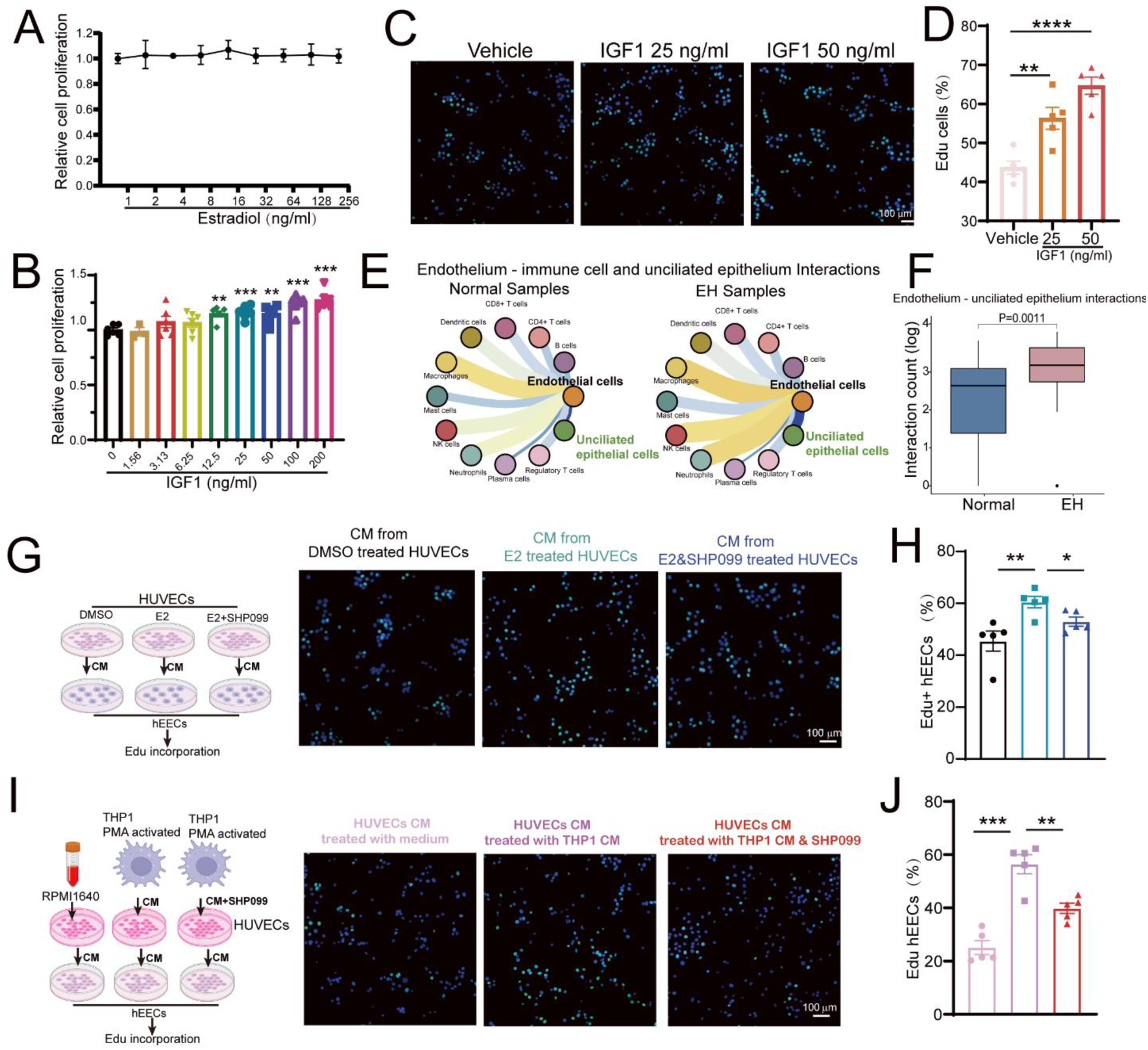
Increased IGF1 secretion by endothelial cell inflammatory activation promoted endometrial epithelial cell proliferation. **A**, CCK8 detection of hEECs proliferation after treatment with different doses of estradiol for 48 h. **B**, CCK8 detection of hEECs proliferation after treatment with different doses of recombinant IGF1 for 48 h. **C**, Edu incorporation after IGF1 stimulation for 6 h and then 20 μM Edu was added into medium for 3 h culture. **D**, The number of Edu^+^ hEECs after different doses of IGF1 stimulation (n=5). **E**, Analysis of cell-cell interaction between endothelial cells and immune cells and epithelial cells by CellPhoneDB using scRNA-seq data from healthy normal endometrium (Normal) and EH endometrium (EH). **F**, Statistic analysis of the cell interactions between endothelial cells and epithelial cells. **G**, Schematic representation of co-culture HUVECs with hEECs. HUVECs first pretreated with DMSO or estradiol (E2, 40 ng/mL) with or without SHP099 (5 μM) for 24 h and then the medium was replaced with complete ECM and cultured for another 24h.The CM was collected to culture hEECs for 12 h and then subjected to Edu incorporation assay. **H**, The number of Edu^+^ hEECs after different doses of IGF1 stimulation (n=5). **I**, Schematic representation of co-culture HUVECs with THP1 cells: The CM from THP1 cells treated with or without PMA (100 ng/mL) for 24 h and then replaced with complete RPMI 1640 medium for another 24 h. The CM from THP1 or complete RPMI 1640 was subjected to treat HUVECs for 24 h. Then, the CM from THP1 CM activated HUVEC was used to culture hEECs for 12 h and then subjected to Edu incorporation assay. **J**, The number of Edu^+^ hEECs after indicated treatment (n=5). Data were analyzed by Dunnett *t* test in Figure 5D, Student’s *t* test in Figure 5F and Dunnett *t* in Figure 5H-J. *p value <0.05; **p value < 0.01; ***p value <0.001; ****p value < 0.0001. See also figure S6.

### 7. SHP2 promotes endothelial cells inflammatory activation through AP-1 activation by dephosphorylating RIPK1 at Y380 site

To study mechanisms of estradiol and SHP2 in estradiol induced endothelial activation, we performed bulk RNA-seq analysis of HUVECs treated with DMSO, estradiol with or without SHP099 and HUVECs treated with estradiol in the condition of SHP2 knockdown (**Fig. 6A and Supplementary Fig. S7A**). We picked up the different expressed genes with opposite expression patterns between cells treated with estradiol alone and SHP099 plus estradiol (**Supplementary Fig. S7B-E**). We noticed that FOS and FOSB mRNA expression changed after estradiol treatment and these changes were reversed by SHP099 or SHP2 knockdown (**Fig. 6B**). FOS and FOSB are members of FOS family, forming AP-1 transcription factor complex with c-Jun. FOS, FOSB, JUN and JUND were found in endothelial cells in our scRNA-Seq data and the transcription factor activity increased in endothelial cells from EH samples (**Fig. 6D**). Target gene analysis of the 4 transcription factors showed that the cytokine, chemokine and the member of NF-kappa B increased in endothelial cells from EH samples, which involved in endothelial cells inflammatory activation (**Fig. 6E**). Moreover, the genes in IGF1 signaling and metabolism such as IRS2, IGS2 and IGF2R were all increased in EH samples (**Fig. 6E**). The phosphorylation of AP-1 member is an indicator of its activation. Thus, we found that estradiol promoted the phosphorylation of c-FOS and c-JUN, while SHP099 decreased the level of phosphorylation of c-FOS and c-JUN (**Fig. 6E**). Furthermore, the inhibitor of AP-1, T5224, could reverse the increased IGF1 expression induced by estradiol (**Fig. 6F**). Besides, T5224 also decreased the mRNA expression of IGF1, ICAM1, SELE and COX2 (**Supplementary Fig. S7F**). Together, our results demonstrated that SHP2 regulated the phosphorylation of transcription factor AP-1. Furthermore, AP-1 positively regulated the inflammatory gene transcription in endothelial cells.

**Figure 6.**
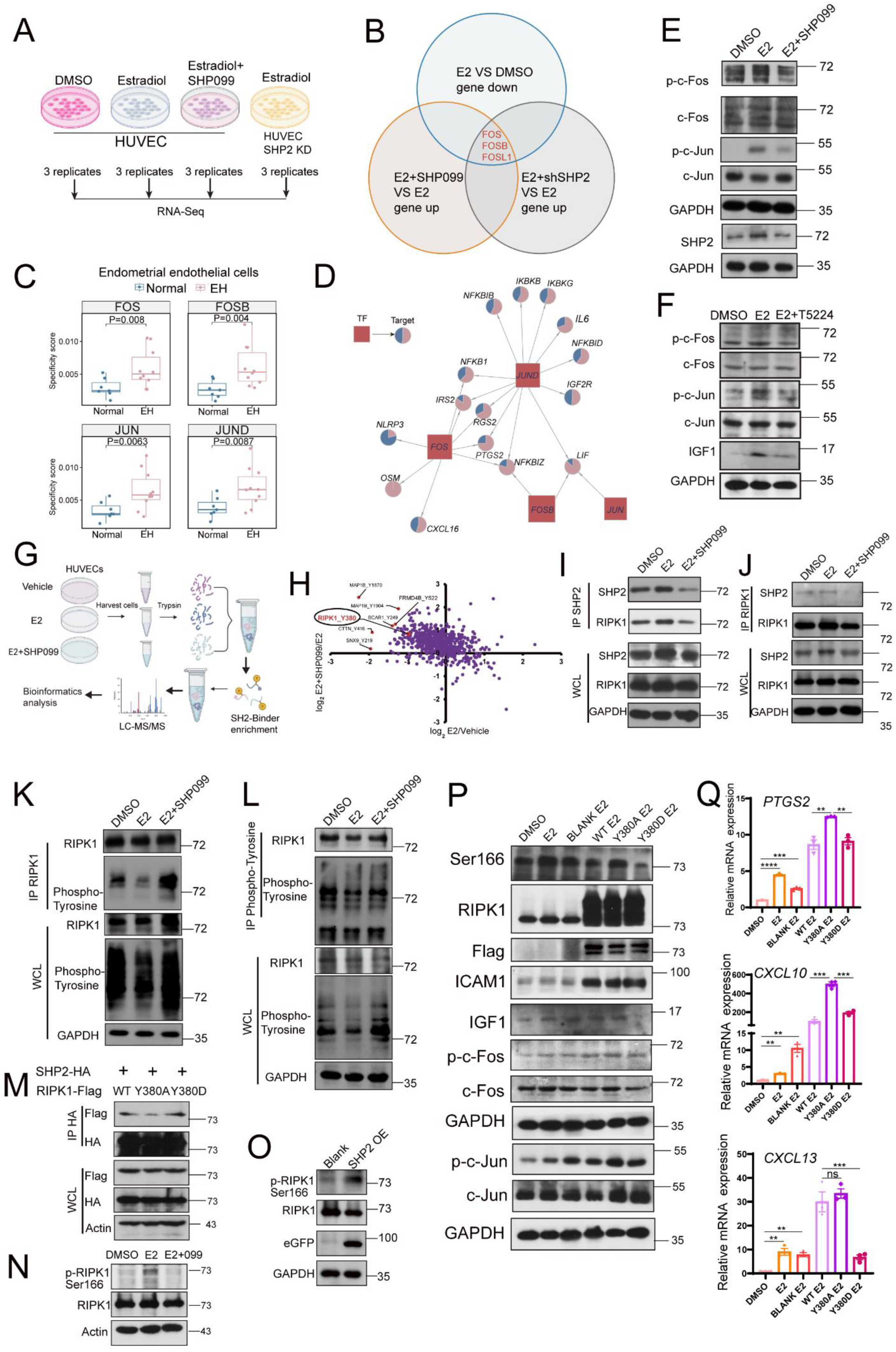
SHP2 promoted IGF1 and inflammatory factors transcription in endothelial cells through RIPK1-AP-1 axis. **A**, Study design of the samples for bulk RNA-Seq (n=3/group). HUVECs were treated with DMSO, estradiol or estradiol plus SHP099 for 6 h and SHP2-knockdown HUVECs treated with estradiol for 6 h were collected and subjected to RNA-Seq for different expressed gene analysis (n=3). **B**, Venn diagram showing the intersection of differentially expressed genes in HUVECs between 2 comparable groups treated as indicated. **C**, The activity of transcription factor, the dot in the box plot represented the activity of indicated transcription factor in each sample. Data are represented as mean± SEM. **D**, Gene regulatory network of transcription factors and corresponding genes. The pie charts showed the different expression of the regulated genes by the indicated gene in normal and EH samples. **E**, WB analysis of AP-1 activation after estradiol (E2) treatment with or without SHP099 (5 μM) for 12 h. **F**, WB analysis of AP-1 activation and IGF1 expression after estradiol (E2) treatment with or without AP-1 inhibitor T5224 pretreated for 3 h. **G**, Experiment outline for stable isotope labeling with amino acids in cell culture-based tyrosine phosphoproteomics in HUVECs treated with vehicle, estradiol (E2, 40 ng/mL) or estradiol (E2, 40 ng/mL) plus SHP099 (5 μM). LC–MS/MS represents liquid chromatography tandem mass spectrometry, n=3/group. **H**, Quadrant diagram for tyrosine phosphoproteomic analysis. E2 represents estradiol. **I** and **J**, Immunoprecipitation (IP) of HUVECs pretreated with SHP099 (5 μM) for 3 hours and then treated with estradiol (E2, 40 ng/mL) for another 6 h. Cell lysates were immunoprecipitated by anti-SHP2 (C) or anti-RIPK1 antibody (D). **K** and **L**, Immunoprecipitation (IP) of HUVECs pretreated with SHP099 (5 μM) for 3 hours and then treated with estradiol (E2, 40 ng/mL) for another 6 h. Cell lysates were then harvested and immunoprecipitated using (E) anti-RIPK1 antibody or (F) anti-phospho-tyrosine antibody for immunoblotting with the indicated antibodies. WCL represents whole cell lysate. **M**, HEK293T cells were transfected with SHP2-HA plasmid and WT-RIPK1 or RIPK1^Y380A^ or RIPK1^Y380D^ point mutant plasmid for 48 h. Then cells were harvested and immunoprecipitated using anti-HA antibody. **N**, WB analysis of the activation of RIPK1 (the level of the phosphorylation of the Ser166 site) after treatment with E2 or E2 plus SHP099. **O**, WB analysis of the activation of RIPK1 after SHP2 overexpression. See also figure S7 and figure S8. **P**, WB analysis of endothelial cell activation related proteins and IGF1 expression after overexpression of RIPK1-WT, RIPK1-Y380A or RIPK1-Y380D by virus in HUVECs. **Q**, Gene expression of endothelial cell activation after overexpression of RIPK1-WT, RIPK1-Y380A or RIPK1-Y380D by virus in HUVECs. Dunnett *t* in Figure 6Q. **p value < 0.01; ***p value <0.001; ****p value < 0.0001.

Because SHP2 is a kind of protein tyrosine phosphatase, we employed quantitative tyrosine phosphoproteomic analysis using mass spectrometry–based stable isotope labeling with amino acids in cell culture (SILAC) to investigate the substrate of SHP2 involved in the regulation of AP-1 phosphorylation (**Fig. 6G**). A total of 1329 phospho-Tyr sites were identified, 934 quantified 26hosphor-Tyr sites included (**Supplementary Fig. S8A**). Considering that SHP2 can dephosphorylate its substrate protein, we got the phosphorylation sites significantly downregulated in estradiol group compared with DMSO group, and also significantly upregulated in estradiol plus SHP099 group compared with estradiol group (**Supplementary Fig. S8B-G**). To find a protein involved in endothelial inflammatory activation and protein phosphorylation[43], we noticed a protein kinase receptor-interacting protein kinase 1 (RIPK1^Y380^) (**Supplementary Fig. S8H**). RIPK1 is a key regulator of inflammation and cell death[44, 45]. The phosphorylation of RIPK1^Y380^ was significantly decreased after estradiol treatment but increased when co-treated with SHP099 (**Fig. 6H**). Therefore, we considered RIPK1 was a substrate of SHP2 and may be responsible for the phosphorylation of c-FOS and c-JUN. To investigate how increased SHP2 regulating the endothelial activation through interacting with RIPK1, we performed coimmunoprecipitation assay to find that the endogenous interaction of SHP2 and RIPK1 was enhanced by estradiol treatment in HUVECs. However, SHP099 disrupted the interaction between SHP2 and RIPK1 (**Fig. 6I, J**). In line with the results of coimmunoprecipitation assay between SHP2 and RIPK1, the tyrosine phosphorylation on RIPK1 decreased after estradiol treatment but enhanced by the SHP099 treatment (**Fig. 6K, L**). All these results indicated that estradiol disrupted the interaction between SHP2 and RIPK1 and decreased the tyrosine phosphorylation of RIPK1. To validate the specific tyrosine site of RIPK1^Y380^, we constructed two RIPK1 mutants by replacing tyrosine at site 380 with aspartic acid (Y380D) to form permanent phosphorylation or with glycine (Y380A) to mimic permanent dephosphorylation, and then transfected plasmids into HEK293T cells. Compared to the WT-RIPK1, the binding of SHP2 and RIPK1^Y380A^ was decreased and it was enhanced when overexpressing RIPK1^Y380D^ (**Fig. 6M**). Many sites on RIPK1, such as Ser25 and Y384, are phosphorylated to inhibit its kinase activity[43, 46]. Therefore, we next investigated how the dephosphorylation of RIPK1^Y380^ by SHP2 affected the RIPK1 kinase activity. Functionally, SHP2 dephosphorylating RIPK1 at Y380 site after estradiol stimulation enhanced the Ser166 phosphorylation of RIPK1 in HUVECs, which represented the enzyme activation of RIPK1 (**Fig. 6N**). Moreover, SHP2 overexpression in HUVECs promoted RIPK1 activation indicated by an increasement in the phosphorylation of RIPK1 at Ser166 (**Fig. 6O**). Furthermore, the permanent dephosphorylation of Y380 (Y380A) led to increasement of RIPK1 activity and inflammatory endothelial activation as shown by increased pSer166 of RIPK1 and its downstream p-c-Fos, p-c-Jun, ICAM1, IGF1 and CXCL10/13 (**Fig. 6P and Q**). While permanent phosphorylation mutant of Y380D had the opposite results. Therefore, all results showed that SHP2 enhanced RIPK1 enzyme activity by dephosphorylating RIPK1^Y380^ and finally promoted the inflammatory activation of endothelial cells. Together, our data showed that SHP2 dephosphorylated RIPK1 at Y380 site after estradiol stimulation, which promoted the enzyme activity of RIPK1 to phosphorylate its downstream c-FOS and c-JUN. The activation of c-Fos and c-Jun promoted the transcription of IGF1 and other inflammatory mediators in activated endothelial cells.

### 8. Pharmacologic targeting inhibition of SHP2 alleviates sterile inflammation and endometrial hyperplasia in mice

Finally, we explored whether pharmacologic targeting SHP2 using its allosteric inhibitor SHP099 could alleviated estradiol-induced EH in mice. Similar to the results in **Fig. 2**, SHP099 treatment significantly reduced the uterine size and weight compared to the EH model group (**Fig. 7A, B**). The H&E staining showed that SHP099 treatment diminished the architectural abnormalities and reduced gland-to-stroma ratio of mice uterus (**Fig. 7C**). The abnormal uterine microvascular was less in SHP099 treated mice. (**Fig.7D**). FACS results and IHC results both demonstrated that the infiltration of macrophage into uterus decreased after SHP099 treatment. (**Supplementary Fig. S9A, B** and **Fig. 7E upper**). The expression of COX2, indicating tissue inflammation, declined after SHP099 treatment, which indicated the resolution of endometrium inflammation (**Fig. 7E bottom)**. The growth factor, IGF1 also increased in EH mice uterus but decreased in the SHP099 treated mice (**Fig. 7E middle)**. Moreover, we also observed that the activation of endothelial cells (CD31^+^ICAM1^+^) and increased RIPK1 activation in mice uterus was inhibited by SHP099 treatment (**Supplementary Fig. S9C-D**). All these results supported that targeting SHP2 could alleviated tissue inflammation and thus the EH progression in mice. Taken together, our study found SHP2 promotes endothelial activation through dephosphorylating RIPK1 at Y380 site, which mediates the macrophage recruitment and subsequent tissue long-term sterile inflammation. The inflammatory environment further activated endothelial cells to secrete increased growth factor IGF1 to promote the epithelial cell proliferation and EH progression in a paracrine manner (**Fig. 7F)**.

**Figure 7.**
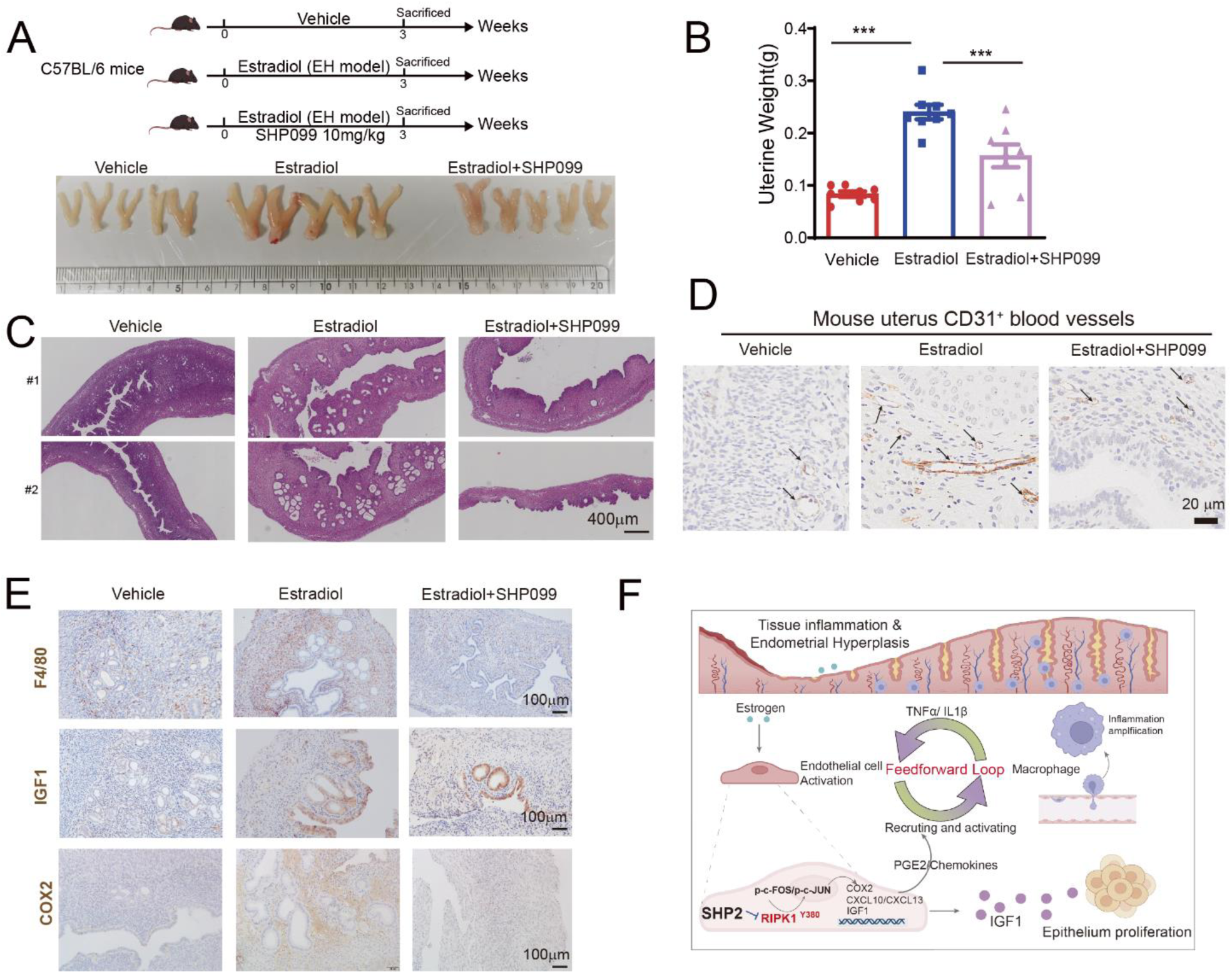
SHP2 inhibition alleviated estradiol induced EH in mice. **A**, Upper, experiment design of 3 groups. Bottom, the morphology of mice uterus from WT mice treated with vehicle (n=8), estradiol (n=8, 100 μg/kg) or estradiol plus SHP099(estradiol, 100 μg/kg; SHP099, 10 mg/kg, n=7) for 21 days. **B**, The uterine weight from mice with indicated treatment. **C**, H&E staining of mice uterus from mice with indicated treatment. **D**, IHC staining of CD31^+^ vessels in mice with indicated treatment. **E**, Upper panel: IHC staining of F4/80^+^ macrophages in mice uterus with indicated treatment. Middle panel: IHC staining of IGF1 level in mice uterus with indicated treatment. Bottom panel: IHC staining of COX2 expression in mice uterus with indicated treatment. **F**, Graphic illustration of the role of SHP2-RIPK1-AP-1 axis in vascular endothelial cell activation mediated tissue sterile inflammation in EH progression. Data are represented as mean ± SEM. Data were analyzed by Tukey-Kramer test. ***p value <0.001. See also figure S9.

## Discussion

Treatment options for EH include sole observation, treatment with progestins and surgery according to the classification of EH, and the choice or physical condition of patients[47]. Although in most cases, progestogen treatment is effective in achieving regression of EH without atypia, there are some women who do not response to and are resistant to either oral or local intrauterine progestogen treatment. The reason for progestogen resistance is complex, but we observed that the estrogen level in the serum of EH patients show no significant difference and is not higher compared with that of healthy individuals. The results indicated that there might be some other factors independent of estrogen that regulate EH diseases. Moreover, some women decline to use progestogen because of the possible side effects[48, 49]. If even a high dose of progestogen is ineffective, disease progression occurs or patients reject to get medical treatment, hysterectomy should be considered as a first-line surgical management. However, the surgical method is not suitable for women who want to preserve their fertility. There is no report about why endometrium inflammation promotes EH and how to find a target for regulating inflammation. Therefore, it is urgent and necessary to further understand the pathological mechanism of tissue inflammation. In our study, we elucidate that endothelial cell activation mediated by increased SHP2 expression has a role in generation of endometrial tissue inflammation and increased IGF1 in the inflammatory environment promotes the abnormal proliferation of endometrial epithelial cells and EH diseases. Therefore, targeting endothelial inflammatory activation through SHP2 inhibition, Y380 phosphorylation of RIPK1 or AP-1, or even using other anti-inflammation targets may be promising strategies for EH medical intervention. Similarly, a recent published study reported that targeting the tyrosine phosphorylation of endothelial VEGFR was a potential anti-inflammatory target[20].

Previous study reported that, genistein, an inhibitor of protein-tyrosine kinases, had an effect on EH treatment[8], which indicated protein tyrosine phosphorylation may be a key regulator in EH. The abnormal expression, enzyme activity alteration, constitutive activation or inactivation mutation of SHP2 occurs in many caner types, inflammatory diseases and reproductive system diseases. Scientists are focusing on finding its inhibitors such as catalytic activity inhibitors PHPS1, allosteric inhibitors SHP099, SHP2 proteolysis targeting chimera molecule SHP2-D26 and molecule that specific targeting the interaction of SHP2 with its substrate[26]. Many kinds of SHP2 inhibitors have gone on clinical trials for the treatment of cancer and inflammatory diseases. Therefore, it is a targetable molecule for medical use. The functions of SHP2 in epithelial cells, immune cells, fibroblasts, chondrocytes and endothelial cells vary due to specific disease conditions. It is widely reported that SHP2 in endothelial cells regulate cell apoptosis[50], destabilization of endothelial junctions through dephosphorylation of VE-cadherin-Y731[51], angiogenesis in tumor[52, 53], recovery of endothelial adherent junctions through controlling of β-catenin phosphorylation[54] and endothelial cell barrier through suppressing VE-cadherin internalization[55]. However, what is the mechanism of SHP2 in endothelial cell activation and how endothelial cell activation mediates tissue inflammation are previously unknown. In our study, we find that estradiol induces the increase of SHP2 in endothelial cells, and consequently trigger cell activation through dephosphorylation of RIPK1 at Y380 site. Previous study reported that dephosphorylation of RIPK1 at Y384 (human, isoform 1) site enhanced the kinase activity[43]. Similarly, our results show that the dephosphorylation of RIPK1(human, isoform 2) at Y380 also promotes the activation of RIPK1 represented as the increased level of phosphorylated Ser166 of RIPK1. Notably, it has been reported that RIPK1 is a key molecule in type Ⅱ endothelial activation[19] and tissue inflammation through different mechanisms[56]. In our study, the activated RIPK1 promotes the activation of transcript factor AP-1, and consequently induce new gene transcription of pro-inflammatory cytokines, chemokines and *IGF1* in HUVECs. The increased cytokines and chemokines reshape the inflammatory environment in endometrium. In line with this, RIPK1 has been previously reported to activate AP-1[57]. However, we do not investigate the precise events that result in the upregulation of SHP2 by estradiol treatment. Besides, we find that the expression of SHP2 in uterine epithelial cell does not change between healthy individuals and EH patients in our scRNA-Seq data. We also confirm that estradiol does not increase the expression of SHP2 in hEECs *in vitro*. Response of SHP2 to estrogen may be different due to cell type and the environment. Interestingly, we unexpectedly find that estradiol alone does not significantly promote the proliferation of hEECs, which is contrary to common knowledge. The surprising results indicate that there are other factors that cooperate with estradiol to promote epithelial cell proliferation.

Since, many research in other groups reported the abnormal activation of endothelial cells may lead to tissue inflammation[19]. Moreover, the association between inflammation and endometrial hyperplasia has been raised for a period of time. Pro-inflammatory cytokines, including TNF-α, IL-1*β* and IL-6, have been observed in EH tissue [58]. Sex steroid hormones are believed to regulate the production and release of cytokine through activating MyD88-dependent TLR- and/or IL-1-receptor signaling pathways[59]. However, how inflammation promotes EH remains unclear. In our study, in line with previous study, we also observe the significantly increased TNF-α, IL-1*β* in the uterus of EH mice model. Considering the important role of immune cell in inflammation, we analyze the interaction between endothelial cells and immune cells. The interaction of endothelial and macrophage is significantly increased in EH patients. So we only pick macrophages and further investigate how macrophages affect EH progression. We find that macrophages are trafficked into the endometrial tissue and activated sequentially by endothelial cells. In turn, activated macrophages enhance the endothelial inflammatory activation in a feedback manner to release more IGF1. The high level of IGF1 thus promotes uterine epithelial cell over-proliferation. However, in our study, we do not investigate how macrophages influence the activation of endothelial cell. But it is reasonable that the cytokines from activated macrophages enhance endothelial cell activation. Besides, the roles of other immune cells in EH, such as neutrophil and T cells, are not clear. Given these, our future studies will be aimed at elucidating the function of other immune cells in EH.

It has been reported that excess estradiol induces simple endometrial hyperplasia which belongs to local hyperestrogenia-mediated diseases. But as tissue inflammation progresses in complex and atypical EH, inflammation in EH is considered a factor in the promotion and progression of pathology, as well as an attributed risk factor for malignancy transformation in EH[2]. It is reported that non-steroidal anti-inflammatory drugs such as COX2 inhibitors reduce the risk of developing certain cancers through regulating cancer-related inflammation[59]. But how tissue inflammation functions in EH remains unknown. In our study, to validate the activation of endothelial cell-generating and sustaining long-term sterile inflammation to promote EH progression, we collected mice uterus at different time points and withdrew estradiol after 3-week EH model construction. Interestingly, the pathology of EH is not regression after estradiol withdrawal because of the existence of tissue inflammation. We further demonstrate that tissue inflammation is the main risk factor for EH progression at the later stage, which is estradiol-independent because of the comparative estradiol level with that of vehicle control mice at the later stage of estradiol withdrawal. Therefore, our study elucidates that it is the continuously activated endothelial cells that sustains the long-time tissue inflammation, which helps activated endothelial cells secret more IGF1 to promote EH progression. Targeting inflammatory mediators (chemokines and cytokines, such as TNF-α and IL-1β), key transcription factors involved in inflammation (such as AP-1), molecules regulating inflammation signaling in inflammatory cells (SHP2, RIPK1) may decrease the incidence and progression of EH.

Taken together, our results demonstrate that high level of SHP2 in endothelial cells promotes cell activation and generates long-term sterile tissue inflammation through trafficking activated tissue macrophages. Consequently, macrophages in the inflammatory endometrium further enhance the continuous inflammatory activation of endothelial cells, independent of continuous estradiol stimulation. Activated endothelial cells release more IGF1, leading to the overproliferation of uterine epithelial cells and the progression of EH. Therefore, our study elucidates the key role of SHP2-mediated endothelial cell activation and chronic sterile tissue inflammation in EH progression. Targeting SHP2 or controlling chronic tissue inflammation through regulating endothelial cell inflammatory activation through RIPK1 may be effective and promising strategies for treating EH.

## Materials Methods

### 1. Mice and endometrial hyperplasia mice model

SHP2^iECKO^ mice were generous gifts from Professor Yuehai Ke (University of Zhejiang, Hangzhou, China). SHP2^iECKO^ mice were generated from SHP2^f/f^ mice being crossed with Cdh5-CreERT2 mice (C57BL/6). SHP2^iECKO^ mice were intraperitoneally injected with tamoxifen (60 mg/kg) for 5 days and left untreated for 7 days to delete endothelial SHP2. SHP2^f/f^ mice were treated with the same dose of tamoxifen as controls. WT C57BL/6 mice were purchased from GemPharmatech Co. Ltd (Nanjing, China). All mice were left free access to standard laboratory chow and water under a condition of 25 °C, suitable humidity, 12 h dark/light cycle. All animal protocols were carried out according to the NIH Guide for the Care and Use of Laboratory Animals and were approved by the Experimental Animal Care and Use Committee of Nanjing University (IACUC-2206005). To establish 17β-estradiol (estradiol, E2) induced endometrial hyperplasia (EH) mice model, female WT mice aged 6-8 weeks or transgenic mice were treated with a daily 100 μg/kg E2 dissolved in olive oil through subcutaneous injection for 21 consecutive days. Mice were sacrificed at 1 day after the final treatment and collected the mice uterus for weight and H&E staining. The severity of endometrial hyperplasia was measured by evaluating the uterine size, uterine weight, architectural abnormalities and gland-to-stroma ratio.

### 2. Cells

HUVECs were isolated from umbilical cord vein by dissociating with 0.2% (w/v) collagenase Ⅱ (1 mg/mL) at 37 ℃ water bath for 20 min with gently rub every 5 min. Then the cells were collected and cultured with ECM (Sciencell, Cat.NO. 1001) supplemented with ECGS. HUVECs between passages 3 and 8 were used in all experiments. Primary lung endothelial cells were isolated from 6-8 weeks-old mice from SHP2^f/f^ or SHP2^iECKO^ mice to confirm the efficiency of SHP2 depletion. Briefly, mice after 5 days tamoxifen injection were anesthetized and subjected to perfusion by intracardiac injection of PBS to remove blood from the lung. Then the lung tissues were cut into pieces and dissociated with a solution containing type-I collagenase (2 mg/mL, Gibco) and DNase (10 μg/mL, Roche) in DPBS for 45 min at 37 °C in a rotatory shaker. After neutralization with DMEM with 20% FBS, the single cell suspension was passed through a 40-μm cell strainer. The CD31^+^ cells were magnetic separation from the cell using anti-CD31 microbeads (Miltenyi, Germany, Cat. 130-097-418). Then the CD31^+^ cells were further stained with anti-mouse CD45 and anti-mouse CD31 and then subjected to FACS sorting to get the CD45^-^CD31^+^ endothelial cells. The human endometrial epithelial cell line (hEEC, WHELAB C1225) was purchased from SHANGHAI WHELAB BIOSCIENCE LIMITED and cultured with MEM supplemented with 10% FBS and 1% NEAA[17]. HEK293T cells and THP1 cells were purchased from ATGG and cultured with DMEM and RPMI1640 medium separately supplemented with 10% FBS and 1% penicillin/streptomycin.

### 3. Clinical samples

The endometrium tissue from EH patients were obtained from EH patients. The detail information of EH patients is list in Supplementary Table 1.

### 4. Single cell dissociation, scRNA-Seq libraries construction and sequencing for endometrium samples

Human endometrial tissues were digested and isolated as previous procedures[17]. Dissociated single cells were then stained with acridine orange/propidium iodide (AO/PI) for viability assessment using a Countstar Fluorescence Cell Analyzer. The proportion of living cells was great than 85%. The scRNA-Seq libraries were generated using the 10X Genomics Chromium Controller Instrument and Chromium Single Cell 3’ V3.1 Reagent Kits. All the libraries were sequenced using an Illumina sequencer (Illumina, San Diego, CA, USA).

### 5. Single-cell sequencing data processing

The 10x Chromium single-cell RNA sequencing (scRNA-seq) data in this study were processed using CellRanger (v3.1.0; 10x Genomics) for read alignment, barcode assignment and unique molecular identifier (UMI) counting, based on the corresponding reference genome (GRCh38). Filtered count matrices from the CellRanger pipeline were converted to sparse matrices using Seurat package (v4.0.0). Cells were filtered with a UMI count greater than 200 and a mitochondrial percentage less than 20%. The ’doubletFinder_v3’ method from the DoubletFinder package (v2.0.3) was applied for additional cell filtering. Filtered data were then log normalized and scaled, with cell–cell variation due to UMI counts and percent mitochondrial reads regressed out. To avoid batch effects among samples and experiments, Seurat’s canonical correlation analysis (CCA) integration tool was used to integrate single-cell data. The top 2000 most variably expressed genes were determined using the “vst” method in the“FindVariableFeatures” function and scaled using “ScaleData” with regression on the proportion of mitochondrial UMIs (mt.percent). Cell clustering was performed by “FindClusters” function at a resolution of 1.5. Dimensionality reduction was performed with “RunUMAP” function and visualized by Uniform Manifold Approximation and Projection (UMAP). For subgroup cell clustering, cells of different types were extracted separately and clustered by their respective top 20 principal components (PCs) using different resolutions based on visual inspection. Marker genes for each cluster were performed using the Wilcoxon rank-sum test (“FindAllMarkers” function with default parameters) and each cell clusters were annotated based on known maker genes the top gene list.

### 6. Pathway analysis

Differentially expressed genes (DEGs) were detected by the ’FindAllMarkers’ function in Seurat. All the gene-set enrichment analyses (GSEA) on DEGs in this study were performed by the clusterProfiler (v3.14.3) package.

### 7. Cell–cell interaction analysis

Cell–cell interactions among the cell types were estimated by CellPhoneDB (v2.1.7) with default parameters (20% of cells expressing the ligand/receptor) and using the version 2.0.0 of the database. CellPhoneDB infers the potential interaction strength between two cell subsets based on the gene expression level of a receptor-receptor pair. The significance of interaction is assessed through a permutation test (1000 times). The normalized gene expression was used as input. Interactions with p value < 0.05 were considered significant. We considered only ligand-receptor interactions based on the annotation from the CellPhoneDB database, and discarded receptor-receptor and other interactions without a clear receptor.

### 8. Gene-regulatory network

To identify cell-type and organ-specific gene regulatory networks, we performed Single-cell Regulatory Network Inference and Clustering (v0.11.2; a Python implementation of SCENIC) in our dataset. Firstly, the original expression data were normalized by dividing the gene count for each cell by the total number of cells in that cell and multiplying by 10,000, followed by a log1p transformation. Next, normalized counts were used to generate the co-expression module with GRNboost2 algorithm implemented in the arboreto package (v0.1.6). Finally, we used pySCENIC with its default parameters to infer co-expression modules using the above-created RcisTarget database. An AUCell value matrix was generated to represent the activity of regulators in each cell. Gene regulatory networks (GRNs) were visualized by the igraph package in R.

### 9. Immunoblotting, immunofluorescence staining and coimmunoprecipitation

For immunoblotting, cells were collected and lysed with WB and IP lysis. Then the extracted proteins were quantified with BCA. Equal quantity of proteins from different samples were separated with SDS-PAGE and transferred onto polyvinylidene difluoride membranes. The cropped membranes were hybridized with different primary antibody at 4 ℃ overnight. Then washing the bands with PBST for 3 times and incubated with secondary antibodies of corresponding species for 1 h. After 3 times washing, the bands were immunoblotting with ECL kit. The primary antibodies used were list in Supplementary Table 2.

For immunofluorescence staining and immunohistochemistry, the tissues were fixed in 4% PFA and embedded in paraffin. 5 mm-thick sections were used. The tissue sections were deparaffinized, rehydrated, and antibody retrieved with sodium citrate. For cell immunofluorescence, the cells were fixed with 4% PFA and permeabilized with 0.5% triton for 15 min. The samples were blocked with 5% goat serum in PBST and incubated overnight at 4 °C with primary antibodies. After 3 times washing, the samples were incubated for 1 h at room temperature with the secondary antibodies (Supplementary Table 2). Then for immunohistochemistry, proteins were detected using the Real Envision Detection kit (GeneTech) according to the manufacturer’s instructions. The nuclei were stained with hematoxylin. For immunofluresence staining, the nuclei were stained with 4′,6-diamidino-2-phenylindole (DAPI) (Beyotime). The samples were then mounted with fluoromount-G (Southern Biotech), and images were acquired using a confocal microscope. The images were further processed using image J software.

For coimmunoprecipitation, the cells were treated accordingly and then lysed with WB or IP lysis and quantified by BCA assay. Equal proteins of 1 mg proteins from each sample were incubated with 2 ug of primary antibody at 4 ℃ overnight and then precipitated with magnetic Protein A/G beads (Millipore) at 4 ℃ for 4 h. Proteins not binding to the beads were washed away by 6 times cold PBS and 2 times cold lysis buffer. Next, the beads were boiled for 10 min in 2X SDS loading buffer. The targeted protein was detected by western blots with the indicated antibodies (Supplementary Table 2).

### 10. SHP2 enzyme activity assay

HUVECs were treated with different doses of estradiol for 48 h and lysed in WB and IP lysis buffer supplemented with PMSF and protein phosphatase inhibitor (Roche) and quantified by BCA assay. Proteins (2 μg) were fixed volume to 100 ul with buffer (60 mM HEPES, pH 7.2, 75 mM NaCl, 75 mM KCl, 1 mM EDTA, 0.05% P-20, and 5 mM DTT) in a 96-well black plate at 25°C for 30 min, and the surrogate substrate DiFMUP (Invitrogen) was added and incubated at 25°C for another 30 min. The fluorescence signal was measured using a microplate reader (Envision) with excitation and emission wavelengths of 340 and 450 nm, respectively. Besides, we set a group with proteins and PHPS1(20 μM, a SHP2 inhibitor) as a control to exclude the effect of all other protein phosphatases. Finally, the subtraction of these two groups was calculated and represented the enzyme activity of SHP2 in HUVECs after estradiol treatment.

### 11. Quantitative PCR

Total RNA was extracted from cells or uterus using Trizol reagent following the manufacturer’s procedure. 1000 ng of RNA were performed reverse transcription to cDNA using Hiscript Ⅱ Q RT SuperMix for qPCR (Cat#: R223, Vazyme Biotech Co,Ltd, China). Quantitative RT-PCR was conducted using Taq Pro Universal SYBR qPCR Master Mix (Cat#: Q712, Vazyme Biotech Co,Ltd, China) on a CFX 100 (Bio-Rad, Hercules, CA) cycler. The primers used were listed in the Supplementary Table 3. Gene expression was calculated using the equation RQ = 2^−△△Ct^ and normalized to *GAPDH* or *ACTB*.

### 12. ELISA assay

Cell culture supernatant was collected and was used to detect IGF1 level by Human IGF-1 ELISA Kit according to the manufacture’s procedure (MULTISCIENCES, EK1131). Mice serums were collected and diluent 2 or 5 times and then were used to detect estradiol level at different time point by QuicKey Pro Mouse E2(Estradiol) ELISA Kit according to the manufacture’s procedure (Elabscience, E-OSEL-0008).

### 13. Monocyte adhesion assay

THP1 cells were first labelled with CM-Dil (2.5 μg/mL) for 10 min at the density of 1X10^6^ cells/mL and then 1 mL of the labelled THP1 cells were incubated with HUVECs with different treatments for 60 min at 37 ◦C. Nonadherent THP1 cells were washed softly with PBS for 3 times. Fluorescence images of adherent THP-1 cells were captured and analyzed using ImageJ software to calculate the number of adherent monocytes.

### 14. CCK8 and Edu incorporation assay

After hEECs were treated with indicated condition for indicated time in a 96-well plate at a volume of 100 ul, 10 ul of CCK8 was added to each well and cultured for another 2 h. Then the absorbance at 450 nm was detected by microplate reader.

For Edu incorporation assay, hEECs were seeded in a 24-well plate at a density of 4X10^4^ per well and cultured overnight and then were subjected to indicated treatment for indicated time. EdU (Beyotime, 20 μM) was added into the medium for 3 h at 37 °C and then fixed in 4% PFA and permeabilized with 0.2% trixton-100. After 3 times washing with PBS, the cells were incubated and protected from light using a click additive solution and nucleus were stained with DAPI. Images were obtained using a fluorescence microscope.

### 15. Cytoplasmic protein and nuclear protein extraction

HUVECs were cultured in 10 cm dishes and treated with estradiol or DMSO for 48 h. Then, cells were collected by trypsin and washed with PBS. The cells were lysed by adding 400 ul of 1% NP40 buffer (10 mM Tris-HCl, pH=7.5) and then the cell lysis was added to 1 mL of the 24% sucrose solution drop by drop and centrifuged at 2500 g for 10 min. Collected the supernatant cytoplasmic protein and the most upper 200 ul of cytoplasmic proteins were used. Then the precipitation was washed with EDTA-PBS solution (0.5 mM) at 4 ℃, 3500 g, 10 s and then washed with PBS for 30 s. The remaining precipitation was cell nucleus. The cell nucleus was lysed with RIPA buffer on ice for 30 min and then centrifuged to get nuclear protein extraction.

### 16. RNA-sequence analysis

Total RNA was extracted using Trizol reagent following the manufacturer’s procedure. The total RNA quantity and purity were analysis of Bioanalyzer 2100 and RNA 6000 Nano LabChip Kit (Agilent, CA, USA, 5067-1511), high-quality RNA samples with RIN number > 7.0 were used to construct sequencing library. Then the purified and cleaved RNA fragments were reverse-transcribed to create the cDNA by SuperScript™ II Reverse Transcriptase (Invitrogen, cat. 1896649, USA). At last, the cDNA libraries were subjected to the 2×150bp paired-end sequencing (PE150) on an Illumina Novaseq™ 6000 following the vendor’s recommended protocol. RNA-seq data analysis was performed by LC-Bio Technology CO., Ltd., Hangzhou, China with the Cloud Analysis Platform (https://www.omicstudio.cn/).

### 17. SILAC and mass spectrometry analysis

HUVECs were cultured in Lys/Arg free DMEM:F12 medium (thermo scientific Cat: 88370) resupplemented with heavy, middle or light Lys and Arg for a minimum of 6 population doublings to make the incorporation of heavy, middle and light isotopes being >90%. Then cells labelled with light, middle and heavy medium were incubated with DMSO, estradiol and combination of estradiol and SHP099 for 6 h, respectively. Washing the cells by cold PBS for three times. Adding 10 cell volumes of 8 M urea lysis buffer (8 M urea, 100 mM NH4HCO3, protease inhibitors and phosphatase inhibitors (Roche) for 30 min on ice. Then sonicating 2s with 5s interval for 3 min with 30% energy. Samples were centrifuged at 12000 rpm for 10 min. The extracted proteins were then digested with trypsin, mixed in equal proportions, and analyzed by mass spectrometry. Postanalysis was done with MaxQuant software.

### 18. Statistics

Graphpad prism 8 software was used for all of the statistical analysis. We assessed data for normal distribution and similar variance between groups. Unpaired two-tailed Student’s *t* test, Dunnett *t* test, Tukey-Kramer, Dunnett were used to assess statistical significance between two groups. One-way analysis of variance (ANOVA) with Tukey’s multiple comparisons was used when comparing three or more groups. *P* value less than 0.05 was considered significant. All data were represented as the mean±SEM.

## Acknowledgments

We thank Professor Yuehai Ke (Zhejiang university) for kindly providing the Shp2^iECKO^ mice. The complementary DNA for Src SH-2 domain triple-mutant T183V/C188A/K206L used in SALIC for the enrichment of tyrosine phosphorylation modification was a gift provided by Precision Proteomics Inc. (Ontario, Canada).

## Author contributions

Conceptualization: Jie Pan, Qiang Xu, Qianming Du, Wen Lv, Yang Sun

Methodology: Jie Pan, Lixin Zhao, Wen Fang, Jiao Qu, Linhui Zhai, Minjia Tan

Investigation: Jie Pan

Visualization: Jie Pan

Writing—original draft: Jie Pan,

Writing—review & editing: Jie Pan, Lixin Zhao, Yang Sun

## Ethics Approval and Consent to Participate

All animal experiments in the present study were undertaken in accordance with the National Institutes of Health Guide for the Care and Use of Laboratory Animals, with the approval of the Animal Care and Use Committee of Nanjing University.

## Data and materials availability

All data are available in the main text or the supplementary materials.

The scRNA-seq data of this study have been deposited into CNGB Sequence Archive (CNSA) of China National GeneBank DataBase (CNGBdb) with accession number: Human: CNP0002449. (http://db.cngb.org/cnsa/project/CNP0002449_b5cf1498/reviewlink/)

In addition, the scRNA-seq data have also been uploaded to Gene Expression Omnibus (GEO) with accession number GSE223643. (https://www.ncbi.nlm.nih.gov/geo/query/acc.cgi?acc=GSE223643)

Bulk RNA-Seq data have been uploaded to GEO with the accession number GSE247027. (https://www.ncbi.nlm.nih.gov/geo/query/acc.cgi?acc=GSE247027)

However, all data remain in private status with reviewer access by mtsfceiqznclhqt of GSE223643 and by gvgrsaykvrsflkr of GSE247027.

---

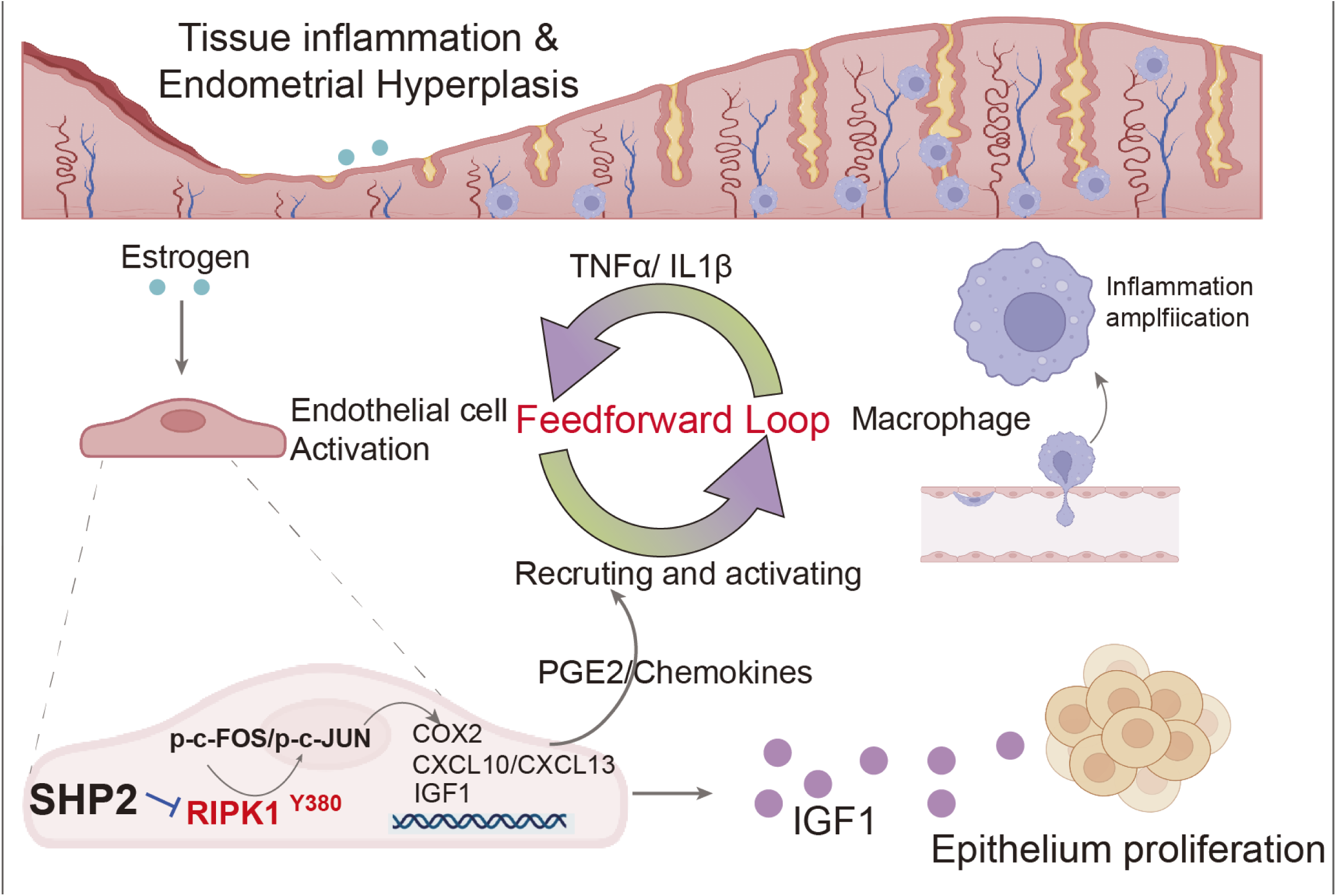

